# An extensive forward genetic screen identifies the HWS-AGO1 axis as pivotal for target mimicry-mediated miRNA degradation in *Arabidopsis*

**DOI:** 10.1101/2025.11.08.687406

**Authors:** Lin Yang, Mengyuan Liu, Mengwei Guo, Youhong Fan, Youchao Wang, Guiting Kang, Ning Li, Yixiao Fang, Jun Mei, Xumin Zhang, Jinxiao Yang, Guodong Ren

**Affiliations:** State Key Laboratory of Genetics and Development of Complex Phenotypes, Zhangjiang mRNA Innovation and Translation Center, School of Life Sciences, Fudan University, Shanghai 200438, China; Beijing Key Laboratory of Maize DNA Fingerprinting and Molecular Breeding, Maize Research Institute, Beijing Academy of Agriculture and Forestry Sciences, Beijing 100089, China; HengJiu Genediting (Beijing) Technology Co., Ltd, Beijing, 100085, China; College of Life Sciences, Zaozhuang University, Zaozhuang 277160, China; Plant Genomics & Molecular Improvement of Colored Fiber Lab, College of Life Sciences and Medicine, Zhejiang Sci-Tech University, Hangzhou 310018, China

**Keywords:** AGO1, miRNA, target mimicry, HWS, degradation

## Abstract

Target RNAs have emerged as key regulators of miRNA stability, thereby influencing development and physiological processes in both plants and animals. The F-box protein HAWAIIAN SKIRT (HWS) has been characterized as a crucial factor in targeted <u>m</u>iRNA <u>d</u>egradation in <u>p</u>lants (pTMD), a process triggered by a specific type of target RNA called target mimicry. However, the precise mechanism by which HWS functions is still poorly understood. We previously established a genetic reporter system based on Short Tandem Target Mimicry of miR160 (STTM160) triggered miR160 degradation, resulting in pleiotropic developmental defects that are easy to monitor. Here, through an extensive forward genetic screen, we identified fourteen additional *hws* alleles that near-completely restored *STTM160*-induced developmental defects, highlighting a central role for HWS in pTMD. Intriguingly, we discovered two adjacent amino acid substitutions (R421K and G422D) in the PAZ domain of AGO1 that significantly suppressed the *STTM160* phenotype. Similar to HWS dysfunction, the AGO1 R421K substitution caused a significant retention of target mimicry RNAs in AGO1 immunoprecipitates, suggesting that the R421 residue of AGO1 may cooperate with HWS for the clearance of AGO1-miRNA-target mimicry complexes. Although the R/G mutations only impacted a small set of endogenous miRNA levels, they were sufficient to rescue the developmental defects caused by *HWS* overexpression (*HWS-OE*), suggesting that the developmental abnormalities caused by *HWS-OE* largely depend on the AGO1-miRNA pathway. Taken together, our findings provide a solid genetic evidence for the coordination of HWS and AGO1 in pTMD.

## Introduction

MicroRNAs (miRNAs) are 21-24 nt non-coding RNAs that orchestrate gene expression at the post-transcriptional level, profoundly influencing plant reproduction, development, and responses to various environmental stimuli (Song et al, 2019). Given their critical biological functions, miRNA dosages are tightly controlled *in vivo*. Abnormalities in overall miRNA expression levels result in severe developmental defects or even lethality (Xu and Chen, 2023). Overexpression or knockdown of a single miRNA can also cause developmental defects and/or changes in stress resistance (Li et al, 2017; Song et al., 2019). miRNA dosages are net effects of biogenesis and degradation. In plants, miRNA biogenesis involves transcription by Pol II, sequential processing by Dicer-like 1 (DCL1), methylation of the 3’ terminus by Hua Enhancer 1 (HEN1), and loading into Argonaute (AGO) proteins to form the RNA-induced silencing complex (RISC) (Yu et al, 2025). Moreover, dozens of proteins have been reported to regulate different steps of the miRNA biogenesis pathway (Li and Yu, 2021; Xu and Chen, 2023). By contrast, the molecular mechanism governing post-biogenesis stability control — particularly degradation mechanisms — is much less understood.

Factors currently known to influence miRNA stability and degradation in plants mainly include the methyltransferase HEN1 (Yu et al, 2005), the terminal nucleotidyl transferases HEN1 suppressor 1 (HESO1) (Ren et al, 2012; Zhao et al, 2012) and UTP: RNA uridylyltransferase 1 (URT1) (Tu et al, 2015; Wang et al, 2015), 3’ to 5’ exonucleases including Small RNA Degrading Nucleases (SDNs) (Chen et al, 2018; Ramachandran and Chen, 2008; Yu et al, 2017) and Atrimmer 2 (ATRM2) (Wang et al, 2018), the effector protein AGO1 (Vaucheret et al, 2004), and target RNAs (Franco-Zorrilla et al, 2007; Wang et al, 2019). HEN1-catalyzed 2’-O-methylation protects the miRNA 3’ end from attack by tailing (mainly HESO1- and URT1-mediated uridylation) and trimming enzymes (Li et al, 2005; Wang et al., 2019; Zhai et al, 2013). AGO1, the primary effector, protects miRNAs from degradation but also recruits decay factors like SDNs, ATRM2, HESO1, and URT1 (Chen et al., 2018; Ren et al, 2014; Wang et al., 2018; Zuber et al, 2018). Notably, a growing body of evidence pinpoints the importance of target RNAs on miRNA stability control. One groundbreaking example is the discovery of the target mimicry long non-coding *RNA Induced by Phosphate Starvation 1* (*IPS1*). *IPS1* harbors a three-nucleotide insertion at the miR399 binding site, forming a central bulge that resists AGO1-mediated cleavage. Under prolonged phosphate-starvation conditions, *IPS1* and its homologous RNAs (*AT4*, *AT4.1*, and *AT4.2*) are strongly induced and sequester miR399, preventing its silencing activity on the *Phosphate 2* (*PHO2*) (Franco-Zorrilla et al., 2007; Shin et al, 2006). Based on the *IPS1* framework, artificial target mimicry technologies have been developed, such as MIM (Target Mimics) (Todesco et al, 2010), STTM (Short Tandem Target Mimics) (Yan et al, 2012) and SPONGE (Reichel et al, 2015). Notably, these synthetic constructs not only suppress miRNA activity but also cause a marked reduction in miRNA abundance, consequently resulting in phenotypic changes (Peng et al, 2018; Reichel et al., 2015; Todesco et al., 2010; Yan et al., 2012; Zhang et al, 2017a). Thus, target mimicry can actively trigger miRNA degradation, rather than merely acting as an miRNA “sponge”. Two independent forward genetic screens using *MIM156* and *STTM160* identified HAWAIIAN SKIRT (HWS) as a key factor in targeted <u>m</u>iRNA <u>d</u>egradation in <u>p</u>lants (pTMD) (Lang et al, 2018; Mei et al, 2019). HWS encodes an F-box protein that interacts with Cullin 1 (CUL1) and multiple *Arabidopsis* SKP1-like (ASK) proteins (Gonzalez-Carranza et al, 2007; Ogura et al, 2008; Zhang et al, 2017b), indicating its role as a substrate adaptor in the S-phase kinase-associated protein 1 (SKP1)-CUL1-F-box protein (SCF) E3 ubiquitin ligase complex (Vierstra, 2009; Zhang et al, 2019). In *hws* mutants, both the miRNAs and its corresponding target mimicry RNAs are readily detected in AGO1 immunoprecipitates, leading to an intriguing hypothesis that HWS may mediate pTMD through the clearance of the non-optimal RISC complex (i.e., the AGO1-miRNA-target mimicry ternary complex) (Mei et al., 2019). However, the detailed underlying mechanism, particularly whether degradation relies on the ubiquitin-26S proteasome system (UPS), remains elusive. Additionally, the *sdn1 sdn2* double knockout mutant significantly suppressed the developmental defects caused by *STTM165* (Yan et al., 2012), suggesting that SDN1/2 may also be involved in pTMD.

In this study, we extended our genetic screen of an ethyl methanesulfonate (EMS)-mutagenized *STTM160* population. Our genetic data unambiguously demonstrated HWS as a central player in pTMD. We further uncovered two adjacent residues (R421 and G422) within the PAZ domain of AGO1 that dictate pTMD efficacy. The AGO1 R/G substitutions partially recapitulated the phenotypic effects of HWS dysfunction, both in morphology and in the regulation of endogenous miRNA levels. Importantly, the developmental defects and miRNA reduction in *HWS*-*OE* plants were rescued by AGO1 R/G substitution mutants, suggesting that *HWS*-mediated phenotypic and molecular changes are mainly modulated through the miRNA pathway. Together, our findings bridge critical gaps in understanding how target mimicry triggers plant miRNA decay and provide key insights into the AGO1-HWS coordination.

## Results

### HWS is a central player in target mimicry-mediated miRNA degradation

We previously established an *STTM160*-triggered miR160 degradation transgene reporter and identified 9 *hws* alleles (*hws-6* to *hws-14*) using a forward genetic approach (Mei et al., 2019; Yang et al, 2022) (Fig. 1a, Fig. S1a). To further dissect the molecular mechanisms of pTMD, we screened an expanded EMS-mutagenized population and recovered 14 additional suppressors that nearly completely restored the developmental aberrations caused by *STTM160*. Sanger sequencing revealed that all these 14 suppressors carried missense or nonsense mutations in *HWS* (Fig. 1a, Fig. S1a), highlighting its central role in pTMD. Importantly, most of the missense mutations were highly conserved across vascular plants (Fig. S1b). We noticed that there was a significant enrichment of mutations in the C-terminal region. One nonsense and seven missense alleles located in this region were analyzed in detail, all of which phenocopied the null *hws* mutants in terms of rescuing the *STTM160* phenotype and miR160 abundance (Fig. 1b-d), demonstrating that the functional importance of this region. According to the current annotation, HWS contains an N-terminal F-box motif (aa 40–85), a putative transmembrane segment (aa 120–140), and a Kelch_2-like motif (aa 290–338), with the C-terminal region (aa 339–411) remaining uncharacterized (Gonzalez-Carranza et al., 2007; Lang et al., 2018). To gain structural insight, we modeled the full-length HWS protein using AlphaFold3 (Abramson et al, 2024). The predicted architecture revealed that the whole C-terminal part of HWS (aa 91-411) exhibited a typical Kelch-repeat β-propeller structure comprising six blades, each consisting of four antiparallel β-sheets (Fig. 1e), consistent with the organization of previously reported Kelch-repeat proteins (Ito et al, 1991; Li et al, 2004).

**Fig. 1.**
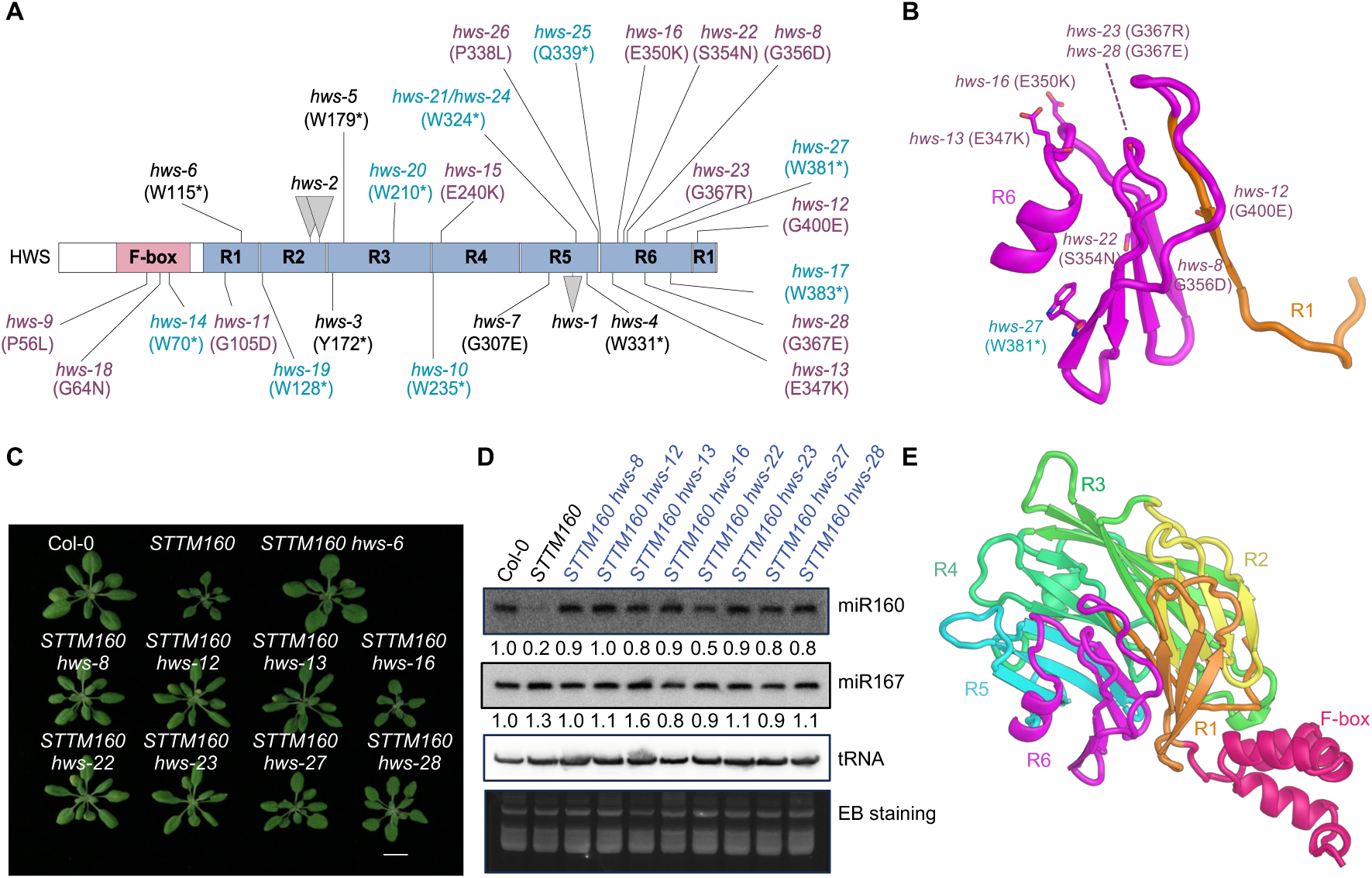
HWS is a central player in pTMD. **(A)** The schematic representation of the HWS protein. Annotated and predicted domains are represented by colored rectangular boxes. Previously published alleles are depicted in black; newly identified mutations are indicated in purple (missense) and blue (nonsense). **(B)** Enlarged view of Kelch-repeat 6 and 1, with mutated residues identified in this region highlighted. The structure was predicted by AlphaFold3. **(C)** Vegetative phenotypes of 3-week-old plants. Scale bar = 1 cm. **(D)** Small RNA Northern blot analysis of miR160 and miR167 in indicated genotypes. Low molecular weight (LMW) RNAs extracted from mixed stages of inflorescence tissues were used. tRNA served as a loading control. **(E)** Predicted 3D structure of HWS by AlphaFold3.

### Two adjacent amino acids in the PAZ domain of AGO1 dictate pTMD efficacy

We also isolated three suppressors (suppressor lines S61, A21, and L38) displaying weaker phenotypic restoration than *hws* (Fig. 2a, Fig. S2a). Sequencing analysis revealed they did not bear mutations in *HWS*, and the *STTM160* transgene remained robustly expressed (Fig. 2b). Small RNA Northern blot analysis showed significant restoration of miR160 expression in all three suppressors, though to a lesser extent than HWS dysfunction (Fig. 2c). We back-crossed the S61 line with *STTM160*, and the F_1_ progeny largely resembled S61. Similarly, the F_1_ progeny of L38 and *STTM160* also resembled L38 (Fig. S2b). These data suggested that the causative mutations in S61 and L38 are dominant. Bulked-segregant analysis coupled with whole-genome sequencing (BSA-seq) mapped the causative mutation to a 7-Mb interval on chromosome 1 containing AGO1 (Fig. 2d). Sequencing data revealed a G to A transition at coding sequence position +1,262 (c.1262G>A) in *AGO1*, resulting in an Arg421 to Lys substitution (R421K). Targeted sequencing of the *AGO1* locus showed that the A21 line carried the identical mutation (i.e., c.1262G>A), while L38 harbored a G to A mutation at +1,265, causing Gly422 to Asp substitution (G422D) (Fig. 2e). Whole-genome re-sequencing indicated that A21 and S61 arose from independent mutagenic events (Fig. S2c, d). For simplicity, we hereafter refer to these alleles by their replaced amino acid (i.e., *AGO1^421K^* and *AGO1^422D^*). Among the ten AGO proteins in *Arabidopsis*, only AGO1 and AGO10 harbor RG residues at the corresponding position, while AGO5 from the same clade contains RT (Fig. S2e). Moreover, the RG residue is highly conserved in plant AGO1 orthologs but is replaced by KG in animals (Fig. S2f). These results indicate that RG-dependent pTMD may be restricted to plant AGO1 and AGO10.

**Fig. 2.**
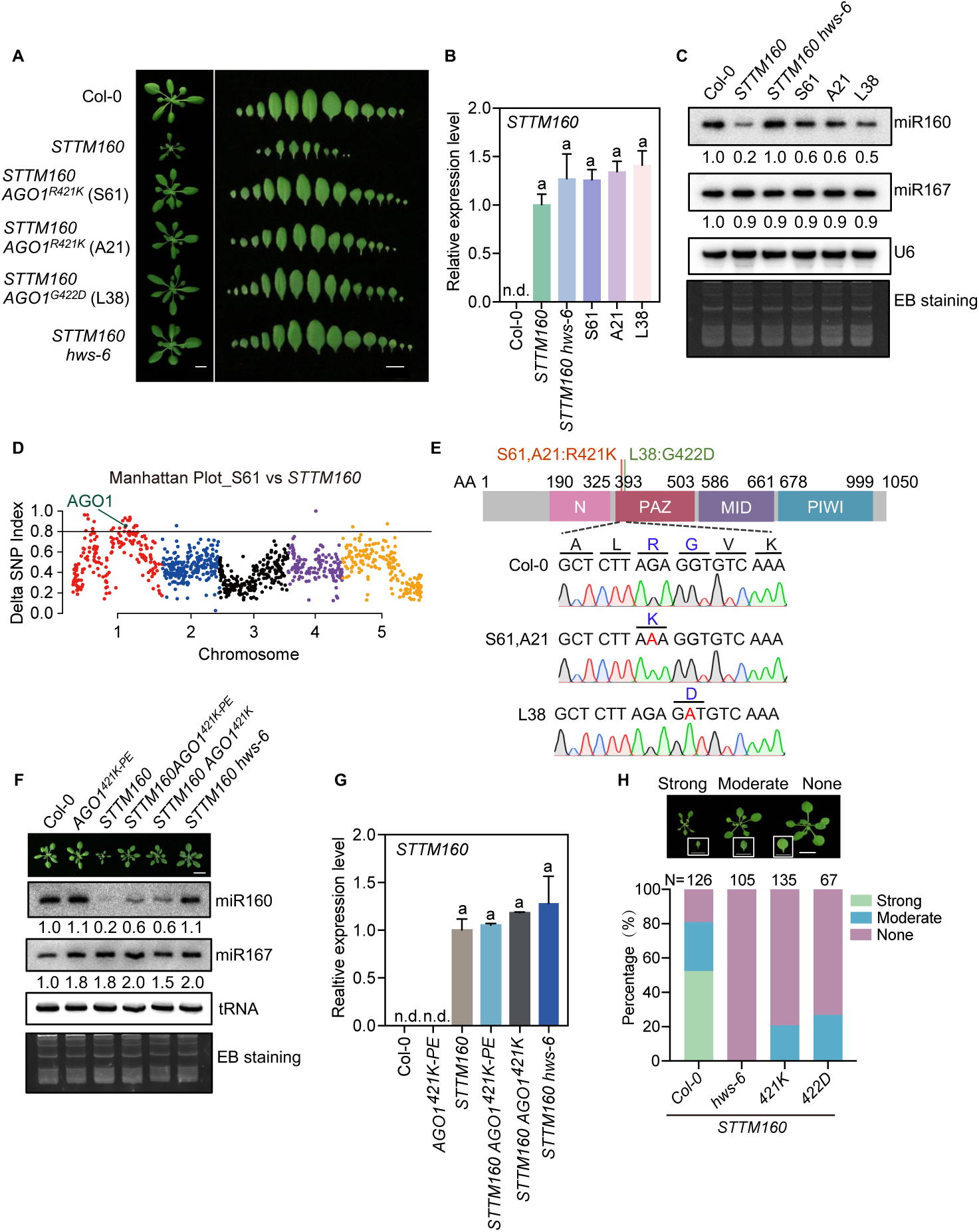
Two adjacent amino acids (R421 and G422) are required for efficient pTMD. **(A)** Vegetative phenotypes of three suppressor lines. The aerial parts of 3-week-old plants were photographed. Scale bar = 1 cm. **(B)** Relative transcript levels of *STTM160* in indicated genotypes quantified by qPCR. *ACTIN2* was used as an internal control. Error bars indicate standard deviation (SD; n = 3 technical replicates). The experiment was repeated twice with similar results. The expression levels in *STTM160* were set to 1. Col-0 was excluded from statistical analysis. No significant differences were observed among the remaining samples (one-way ANOVA, p < 0.05). **(C)** Small RNA northern blot analysis of miR160 and miR167 in indicated genotypes. Low molecular weight (LMW) RNAs extracted from mixed stages of inflorescence tissues were used. U6 and ethidium bromide (EB)-stained RNAs served as loading controls. **(D)** Manhattan plot of the ΔSNP index between the S61-like F_2_ pool and the *STTM160-like* F_2_ pool, with the SNP locus in AGO1 highlighted in green. **(E)** Schematic diagram of the AGO1 domain organization and Sanger sequencing validation of AGO1 genomic mutations. **(F)** *AGO1^PE-421K^* partially suppressed the *STTM160* phenotype. Low molecular weight (LMW) RNAs extracted from 7-day-old seedlings were used for northern blot analysis. tRNA served as an internal control. **(G)** Expression levels of the *STTM160* transcript in different genotypes by qPCR. *UBQ5* served as an internal control. Data represent means ± SD (n = 2 biological replicates). Col-0 and *AGO1^421K-PE^* were excluded from statistical analysis (one-way ANOVA, p < 0.05). **(H)** *AGO1^421K^* and *AGO1^422D^*partially suppressed *STTM160*-induced developmental defects. The *STTM160* construct was transformed into different genotypes, and T_1_ transgenic plants were classified into three categories based on the severity of leaf serration. Numbers shown on the top of the bar graph indicate the number of T_1_ transgenes used for analysis.

Transforming the *AGO1^421K^* construct into the *STTM160* reporter line frequently caused transgene silencing, making it difficult to verify if *AGO1^421K^* was the causal mutation. We therefore switched to using the CRISPR-Cas9 prime editing (PE) technique (Xu et al, 2022) to recreate the R421K allele (*AGO1^421K-PE^*) and segregated out the PE transgene after successful editing (Fig. S3). The *AGO1^421K-PE^* plants were indistinguishable from the wild type during vegetative growth (Fig. 2f). When introduced into *STTM160*, *AGO1^421K-PE^* significantly suppressed *STTM160*-induced miR160 degradation and developmental defects without affecting *STTM160* expression (Fig. 2f-g).

Considering the pTMD sensitivity may be affected by the target mimicry expression levels, we obtained *hws-6* (Mei et al., 2019), *AGO1^421K^*, and *AGO1^422D^*by segregating the *STTM160* transgene away from the F_2_ progenies of the respective back-crossing populations (Fig. 3a) and re-transformed the *STTM160* construct into wild type and the above mutant. We classified *STTM160-*induced developmental defects into three levels (strong, moderate, and none). As shown in Figure 2h, nearly 81% (102/126) of the T_1_ transgenic individuals in Col-0 exhibited typical *STTM160-like* phenotypes (52% strong + 29% moderate). In sharp contrast, no T_1_ individuals in *hws-6* (0/105) showed *STTM160-like* phenotypes, validating HWS’s central role in pTMD. Although no strong *STTM160-like* phenotype was observed in T_1_ individuals of the *AGO1^421K^* and *AGO1^422D^* backgrounds, a significant proportion [21% (28/135) in *AGO1^421K^* and 27% (18/67) in *AGO1^422D^*, respectively] displayed a moderate phenotype, indicating the R/G residues of AGO1 significantly contribute to, but are not essential for, pTMD.

**Fig. 3.**
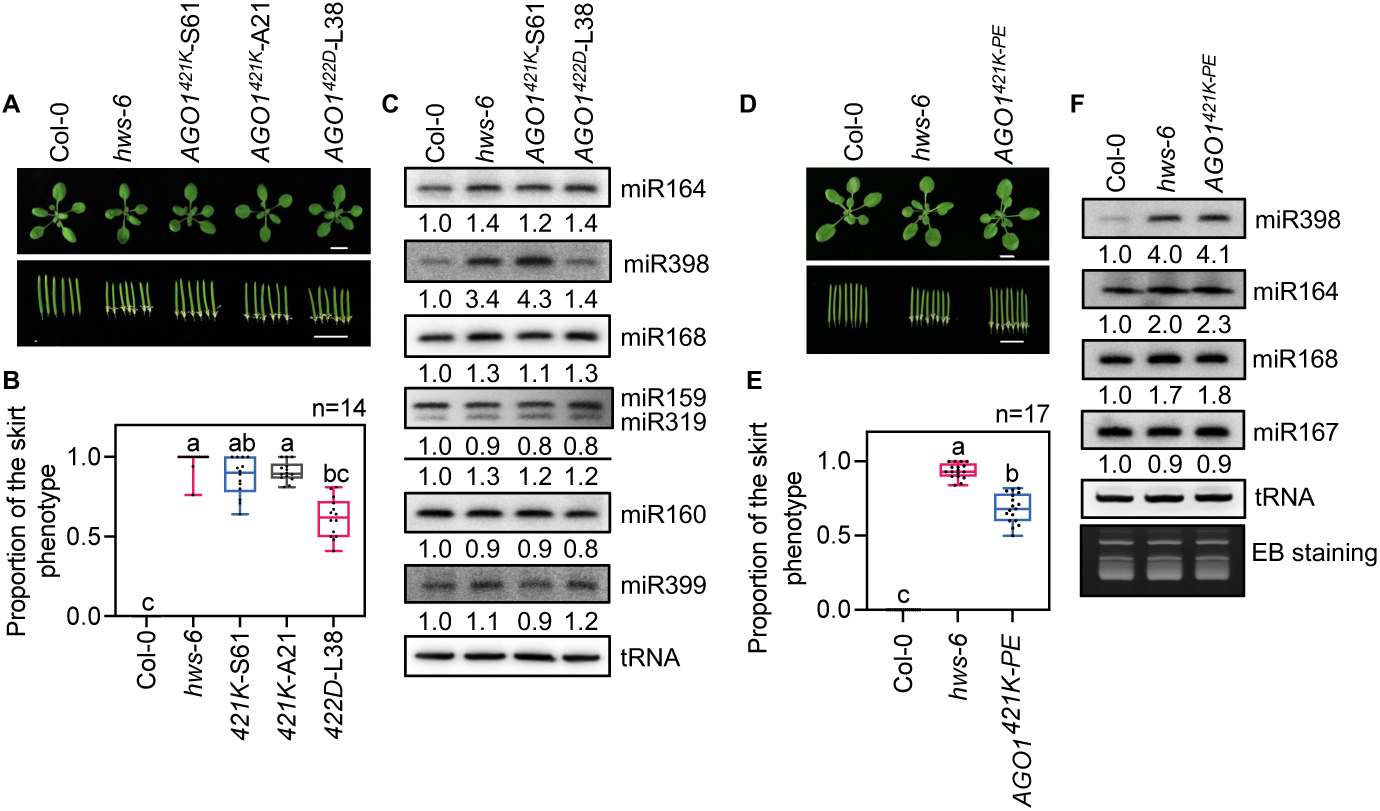
*AGO1^421K^* and *AGO1^422D^* selectively affect a subset of miRNAs. **(A, D)** Vegetative and silique phenotypes of different genotypes. Aerial parts of 18-day-old plants (upper panel) and the sepal-fusion “skirt” phenotype (lower panel) were shown. Scale bar = 1 cm. **(B, E)** Quantification of the “skirt” phenotype in different genotypes. Each dot represents the proportion of siliques with “skirt” on the main stem of one plant. A total of 14 (**B**) and 17 (**E**) plants for each genotype were analyzed. Statistical analyses were performed using the Kruskal-Wallis test with Dunn’s post hoc comparisons. Different letters indicate statistically significant differences among genotypes. **(C, F)** Northern blot analysis of mature miRNA levels in inflorescence tissues of different genotypes. tRNA served as a loading control.

### AGO1^421K^ and AGO1^422D^ selectively perturb endogenous miRNA homeostasis

Similar to *AGO1^421K-PE^*, neither *AGO1^421K^*nor *AGO1^422D^* displayed the severe developmental defects characteristic of *ago1* hypomorphic alleles, indicating that the target silencing activity remains largely intact in *AGO1^421K^* and *AGO1^422D^* (Fig. 2f, Fig. 3a). Nevertheless, both *AGO1^421K^*and *AGO1^422D^* exhibited the “skirt” phenotype (i.e., persistent sepal adhesion on mature siliques), albeit milder than *hws* mutants (Fig. 3a-b). Small-RNA northern blot analysis revealed specific over-accumulation of a subset of endogenous miRNAs (e.g., miR164, miR168, miR398 and miR319) in *AGO1^421K^* and *AGO1^422D^*, as in *hws-6* (Mei et al., 2019), while other tested miRNAs remained largely unaffected (Fig. 3c). Similar results were obtained in *AGO1^421K-PE^* (Fig. 3d-f). Together, these findings suggested that AGO1 and HWS may cooperate to regulate the homeostasis of a discrete miRNA subset.

### *AGO1^421K^* and *AGO1^422D^* suppressed *HWS*-overexpression-induced developmental defects

Although loss-of-function in *HWS* causes only subtle morphological alterations in *Arabidopsis,* its over-expression (*HWS-OE*) results in pleiotropic developmental defects, including dwarfism, serrated leaves, and reduced fertility, concomitant with reduced accumulation of dozens of miRNAs (Gonzalo et al, 2025; Lang et al., 2018; Mei et al., 2019; Zhang et al., 2017b). Yet, the extent to which the developmental defects of *HWS-OE* transgenic plants are attributable to dysregulated miRNA dosages remains unknown. We introduced *AGO1^421K^* or *AGO1^422D^* into *HWS-OE* and found that both *AGO1* allelic mutations suppressed the morphological symptoms of *HWS-OE* in a dominant manner, without significant changes in *HWS* expression (Fig. 4a, c, Fig. S4). Accordingly, the reduced expression levels of miR398, miR164, and miR168 in *HWS-OE* were significantly restored in *HWS-OE AGO1^421K^* and *HWS-OE AGO1^422D^*(Fig. 4b). These results suggested that the developmental defects of *HWS-OE* in *Arabidopsis* may be primarily due to accelerated miRNA degradation.

**Fig. 4.**
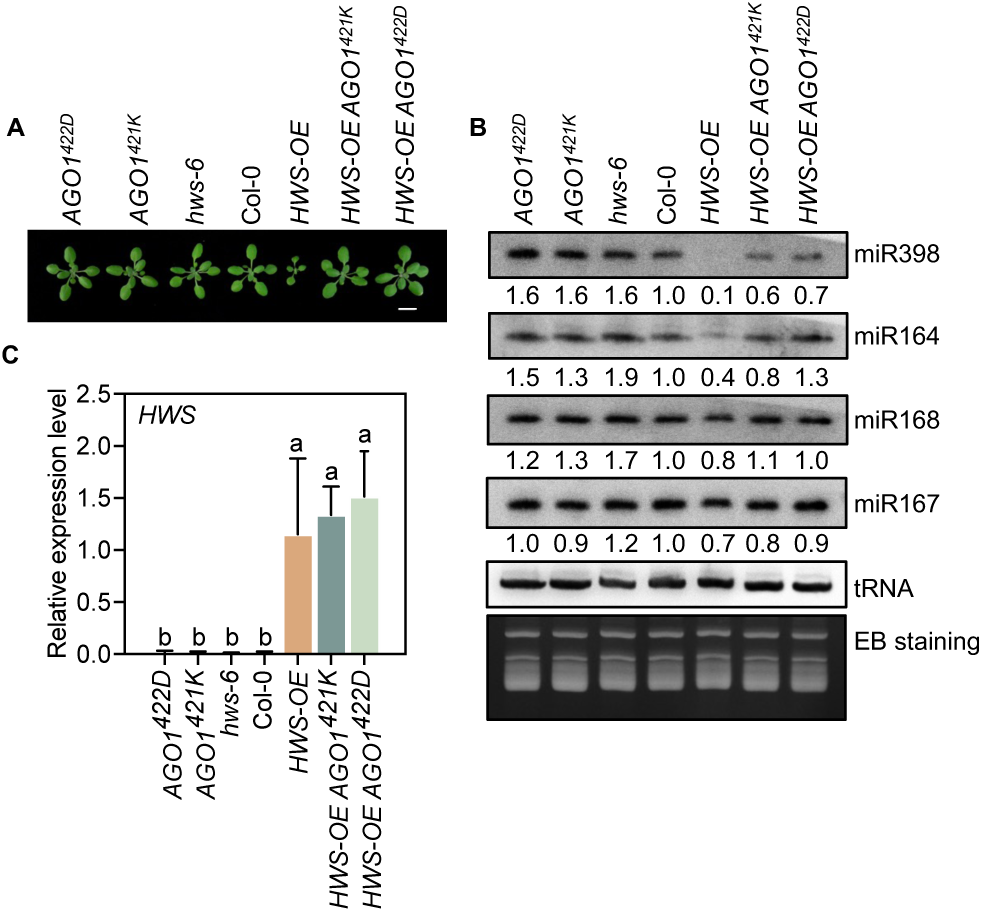
*AGO1^421K^* and *AGO1^422D^* rescue *HWS-OE* induced developmental defects and miRNA reduction. **(A)** Vegetative phenotypes of three-week-old plants. Scale bar = 1 cm. **(B)** Small RNA Northern blot analysis of miRNA levels in inflorescence tissues of different genotypes. tRNA served as a loading control. **(C)** Relative expression levels of *HWS* in different genotypes. *UBQ5* was used as an internal control. Data are presented as means ± SD (n = 3 biological replicates). Statistical analyses were performed using one-way ANOVA. Different letters indicate statistically significant differences among genotypes.

### Retention of non-optimal RISC in *AGO1^421K^*

In *STTM160 AGO1^421K^* and *STTM160 AGO1^422D^*, *STTM160* transcripts remained highly expressed while miR160 levels were significantly restored (Fig. 2b-c), implying that the R/G mutations permit a stable coexistence of target mimicry RNAs and their corresponding miRNAs. We previously reported that HWS dysfunction results in the retention of non-optimal RISC (i.e., the AGO1-miRNA-target mimicry ternary complex) (Mei et al., 2019). To test whether target mimicry RNAs and their corresponding miRNAs coexist in *AGO1^421K^*, phosphorus-starved seedlings from Col-0, *STTM160,* and *STTM160 AGO1^421K^*were subjected to AGO1 RNA immunoprecipitation (RIP) analysis (Fig. 5a). As expected, miR160 was detected in input and AGO1-IP samples of Col-0 and *STTM160 AGO1^421K^*, but not in those of *STTM160*. Interestingly, while the *STTM160* transcript was readily detected in input samples of *STTM160* and *STTM160 AGO1^421K^*, it was only detected in the AGO1-IP sample of *STTM160 AGO1^421K^* (Fig. 5b). These results suggested the specific retention of the “AGO1*^421K^-*miR160-*STTM160*” non-optimal RISC. We also examined the accumulation of miR399 and its endogenous target mimicry transcripts (i.e., *IPS1*, *AT4,* and *AT4.1*) and found that they were enriched in the AGO1-IP sample from *STTM160 AGO1^421K^* (Fig. 5b). The co-existence of miR160, miR399, and their respective target mimicry RNAs in the *AGO1^421K^* immunoprecipitates suggested that the R421 residue of AGO1 may play a crucial role in HWS-dependent non-optimal RISC clearance.

**Fig. 5.**
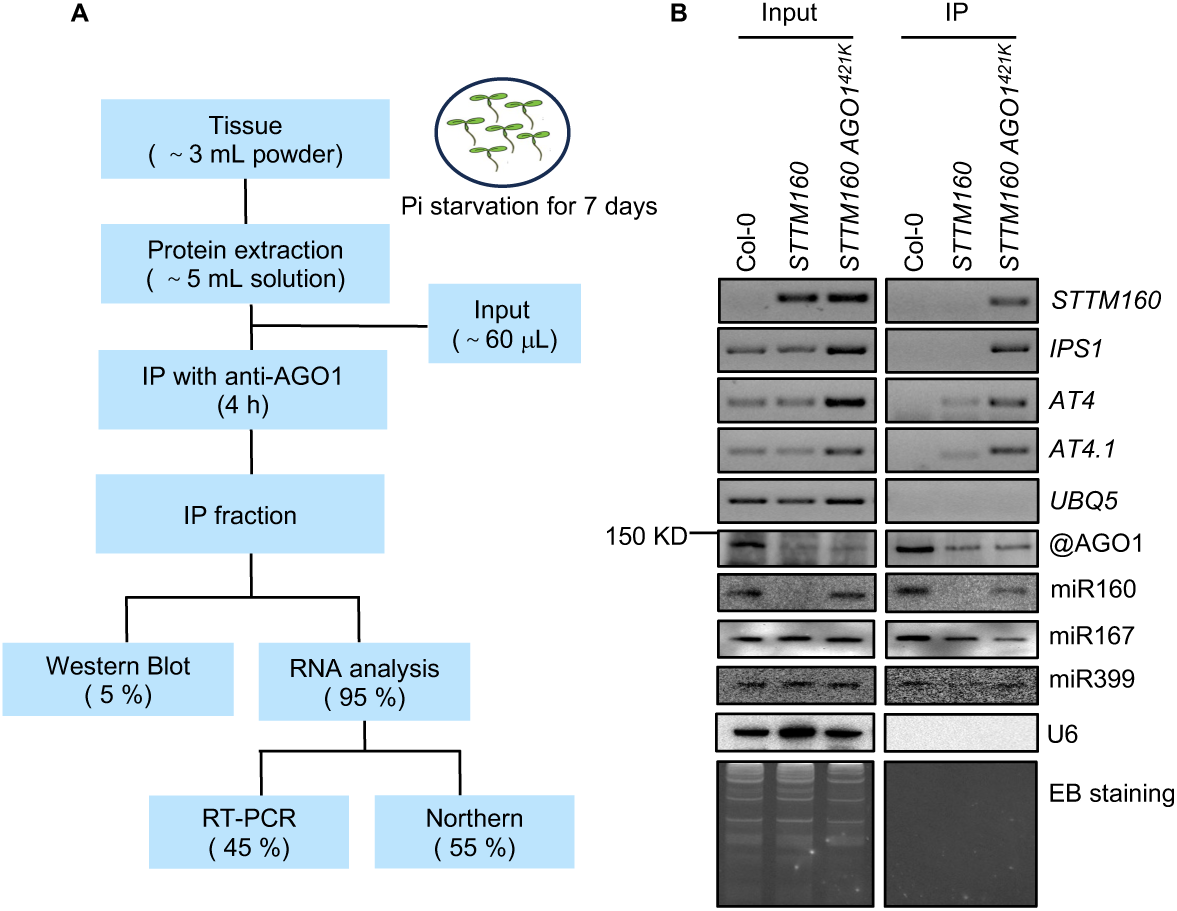
*AGO1^421K^* is required for efficient non-optimal RISC clearance. **(A)** Workflow of the experiment. IP, immunoprecipitation. **(B)** Analysis of target mimicry RNAs and their corresponding miRNAs in AGO1 immunoprecipitates. Target mimicry RNAs were analyzed by reverse transcription PCR (RT-PCR). *UBQ5* served as a negative control. miRNAs were analyzed by Northern blot. U6 and EB staining were used as controls. AGO1 was detected by Western blot.

## Discussion

Our study demonstrates that HWS functions as a core component in mediating pTMD. Mutations arising from forward genetic screens span the Kelch-repeat and the F-box domains, suggesting that both domains are crucial for HWS function. Furthermore, we identified two adjacent residues (R421 and G422) in the AGO1 PAZ domain critical for pTMD efficacy. Importantly, *HWS*-*OE*-induced developmental defects were suppressed by AGO1 R/G substitution mutants. These findings establish a solid genetic link between HWS and AGO1, advancing our understanding of how plants achieve precise control over miRNA homeostasis and dynamic regulation through a specialized degradation pathway.

The role of HWS as an E3 ligase adapter implies its direct recognition of AGO1 conformation and/or post-translational modification changes upon target mimicry binding (Lang et al., 2018; Mei et al., 2019). We propose the following working model: when miRNAs engage target mimicries with central bulges, the non-optimal RISC may expose degrons or protein-protein interaction interfaces recognized by HWS, which recruits ubiquitin machinery to mark AGO1 or associated components for proteasomal degradation. This model aligns with co-immunoprecipitation data showing that target mimicries co-purify with AGO1 in *hws* and *AGO1^421K^* mutants (Fig. 5b), indicating that the HWS-AGO1 axis may be crucial for non-optimal RISC disassembly. Consistent with this hypothesis, previous data showed that HWS interacts with the core SCF component Cullin 1 (CUL1) and multiple ASK proteins (Gonzalez-Carranza et al., 2007; Ogura et al., 2008; Zhang et al., 2017b). The F-box domain is essential for HWS function (Lang et al., 2018; Mei et al., 2019). Moreover, we identified a number of mutations in the Kelch-repeat domain (Fig. 1; Fig. S1), which is known to confer substrate specificity through protein-protein interaction (Hassan et al, 2015). Unfortunately, multiple attempts to detect a robust HWS-AGO1 interaction were largely unsuccessful (Lang et al., 2018; Mei et al., 2019). In this study, we also conducted split luciferase analysis through an agrobacterium-mediated transient expression in tobacco leaves. As a positive control, we validated the F-box-dependent interaction between HWS and ASK1. However, no signals were detected between AGO1-nLUC and cLUC-HWS regardless of co-infiltration of *MIM159* or not (Fig. S5a, b). Thus, while cumulative data pinpoint that HWS and AGO1 likely act closely to modulate targeted miRNA degradation, evidence for this working model remains flimsy. Hitherto, it is equally possible that HWS could modulate pTMD in a non-SCF manner (Liu et al, 2011; Rieu et al, 2023a; Rieu et al, 2023b). Notably, a recent study showed that HWS associates with the nuclear pore complex (NPC) and Mediator subunits and antagonizes Hasty (HST)-mediated *MIRNA* transcription and non-cell-autonomous miRNA movement (Gonzalo et al., 2025). Given that HWS localizes both in the nucleus and cytoplasm (Gonzalo et al., 2025; Lang et al., 2018), it appears that HWS may govern miRNA expression and/or RISC state at multiple steps by targeting different substrates. Further efforts are needed to dissect the distinct contributions and their underlying mechanisms of specific HWS subcellular pools to the miRNA pathway.

Notably, amino acid substitutions in the RG motif of AGO1 (i.e., R421K and G422D) largely suppress pTMD but exhibit weaker phenotypic effects compared to *HWS* knockout. This phenotypic discrepancy suggests the RG motif serves as a regulatory hub rather than an essential degradation signal. How does the RG motif impact pTMD? There are several mechanistic possibilities that are not mutually exclusive. First, the RG di-residue may form a conformational sensor that is exposed upon target mimicry RNA binding (Sheu-Gruttadauria et al, 2019). In this model, arginine’s unique capacity for bidentate hydrogen bonding with phosphate groups (or acidic residues) (Rapp et al, 2013) would enable HWS recruitment. Substitution to lysine (R421K) may abolish such interaction due to lysine’s flexible but monodentate amine group, while G422D introduces a bulky, negatively charged amino acid that may rigidify the local backbone via steric clash and/or electrostatic repulsion (Guan et al, 2016). Second, arginine residues are hotspots for post-translational modifications (PTMs) (Ahmad and Cao, 2012). We speculate that target mimicry binding may induce R421 modification (e.g., methylation by PRMTs), converting the RG motif into a “degron code” recognized by HWS. Substitution with lysine abolishes arginine-specific PTM (Yang et al, 2015), and G422D may sterically block modifying enzymes (Guan et al., 2016). Interestingly, a recent study reported that AGO1 can be symmetrically dimethylated (sDMA) by the type II PRMT5 and asymmetrically dimethylated (aDMA) by yet uncharacterized type I PRMTs (Barre-Villeneuve et al, 2024). Additionally, PRMT5-mediated sDMA of AGO2 is recognized by Tudor-domain proteins TSNs, thereby facilitating proteasome-dependent degradation of AGO2 in *Arabidopsis* (Hu et al, 2019). It would be intriguing to test whether the R421 residue is methylated upon target mimicry binding. Third, the RG motif may facilitate allosteric communication between the PAZ domain and AGO1’s PIWI lobe upon target mimicry binding (Sheu-Gruttadauria et al., 2019). Disruption of this relay (via R421K or G422D) may specifically block the propagation of mimicry-induced structural signals to HWS recognition. Critically, the dominance of R421K/G422D mutations implies that mutant AGO1 competes with wild-type AGO1 for HWS binding but cannot trigger degradation, which sequesters HWS in non-optimal RISCs and attenuates pTMD efficacy. Clearly, all these intriguing possibilities await future investigation.

Despite mild phenotypes in *Arabidopsis*, HWS plays profound physiological roles in crops. Tomato *hws* mutants exhibit facultative parthenocarpy, elevated sugar content, and improved fruit set under warm conditions—traits potentially linked to altered levels of miRNAs such as miR156/miR164 (Damayanti et al, 2019; Lombardo et al, 2021). Rice EP3 and FBK1 (both are HWS orthologs) regulate panicle architecture, grain size, floral organ identity, and root architecture (Borah and Khurana, 2018; Borna et al, 2022; Li et al, 2011; Piao et al, 2009; Yu et al, 2015). Such pleiotropic effects demonstrate the importance of HWS, depending or not on miRNA. Significantly, AGO1 R/G substitution mutants were able to greatly rescue the *HWS-OE* defects in *Arabidopsis*. Thus, it will be intriguing to test whether pleiotropic phenotypes caused by *HWS* malfunction in rice and tomato also depend on the pTMD pathway. Dissection of respective endogenous target mimicries (eTMs) and their controlled physiological traits will be crucial to understanding the physiological significance of pTMD in diverse plant species.

Last but not least, despite functional parallels in triggering miRNA degradation upon central bulged/mismatched target engagement, *HWS* (plants) (Lang et al., 2018; Mei et al., 2019) and *ZSWIM8* (animals) (Han et al, 2020; Shi et al, 2020) represent stark examples of non-homologous innovation. Different from HWS, ZSWIM8 operates as a substrate receptor for Cullin3-RING ligases (CRL3), relying on a distinctive SWIM-type zinc finger domain (Han et al., 2020; Shi et al., 2020). Moreover, the critical RG motif in plant AGO1 is replaced by KG in metazoan AGO2 (Fig. S2f). Together, these key discrepancies imply independent evolutionary origins and different mechanisms with convergent functional output (i.e., targeted miRNA degradation) in plants and animals.

## MATERIALS AND METHODS

### Plant material and growth conditions

*Arabidopsis thaliana* (ecotype Col-0) and *Nicotiana benthamiana* were used in this study. *STTM160*, *STTM160 hws-6*, and *hws-6* were described previously (Mei et al., 2019). For *HWS-OE*, a single-copied line exhibiting pronounced developmental defects was screened from independent T_1_ lines generated previously (Mei et al., 2019). Other mutants were isolated from forward genetic screens or generated via prime editing (see below for details). Plants were grown on half-strength MS medium or in soil under long-day conditions (16 h light/8 h dark, 22°C, 75% relative humidity, ∼120 μmol photons m⁻² s⁻¹) in controlled growth chambers. For phosphate starvation assays, 7-day-old seedlings grown on half-strength Murashige and Skoog (MS) agar medium were transferred to phosphate-deficient solid medium (Coolaber, PM1011-P) for 7 days.

### Forward genetic screen and BSA-seq analysis

Pooled M_2_ populations (each pool contains 15-30 independent lines) of EMS-mutagenized *STTM160* were used for suppressor screen (Mei et al., 2019). 34 F_2_ individuals showing suppressed morphological phenotype and restored miR160 expression from the cross between S61 and *STTM160* were pooled and subjected to Illumina sequencing. BSA-seq and fine-mapping were conducted as previously described (Abe et al, 2012; Mei et al., 2019). Primers used for fine-mapping are listed in Table S1.

### Prime editing

A CRISPR-based prime editing (PE) system was constructed in a binary vector comprising a Cas9 (H840A) nickase fused to Moloney murine leukemia virus (MMLV) reverse transcriptase, flanked by bipartite nuclear localization signals, and a CaMV 35S-driven DsRed reporter. A pegRNA covering the *AGO1^421K^* corresponding genomic region was designed and cloned into the backbone vector pCambia2300 (Xu et al., 2022). T_2_ individuals without Dsred fluorescence (i.e., Cas9 free) were screened for successful PE editing. The primers used for genotyping of edited loci are listed in Table S1.

### Western blot assay

Western blot analysis was carried out following established protocols (Wang et al., 2018). Total protein was extracted from *A. thaliana* seedlings, resolved by SDS-PAGE, and then transferred onto PVDF membranes (Millipore, USA). Immunodetection was performed using a custom-made anti-AGO1 antibody (GL Biochem (Shanghai) Ltd), targeted against the peptide sequence MVRKRRRTDAPS. Anti-HSC70 (Agrisera, AS08371) was used as a loading control. The immunoreactive bands were visualized using lumiQ ECL electrochemiluminescence (ShareBio, China).

### RNA extraction and expression analysis

Total RNA was extracted from *Arabidopsis* tissues using RNAiso Plus reagent (Takara, Code No. 9109) according to the manufacturer’s instructions. Low molecular weight RNAs were enriched and resolved on 16% denaturing polyacrylamide gel containing 7 M urea. Northern blotting was performed with biotin- or ^32^P-labeled oligonucleotide probes following established protocols (Wang et al., 2018). Radioactive signals were visualized with a Typhoon FLA-9000 scanner (GE Healthcare) and quantified using ImageJ. For gene expression analysis, 1 μg of total RNA was reverse transcribed using the HiScript II Q RT SuperMix (+gDNA wiper, Vazyme), and quantitative PCR (qPCR) was performed on a CFX96 detection system (Bio-Rad) with *Ubiquitin* 5 (*UBQ5*) or *Actin2* (*ACT2*) as reference genes. All primers and oligonucleotides used are listed in Table S1.

### Phylogenetic and sequence analyses

Protein sequences for *A. thaliana* AGO proteins, AGO1 orthologs, and HWS homologs across other species were obtained from NCBI (https://www.ncbi.nlm.nih.gov/) and Uniprot (https://www.uniprot.org/uniprotkb) (Su et al, 2024). Multiple sequence alignments were generated in MEGA11 using the ClustalW algorithm. Phylogenetic relationships were inferred using the neighbor-joining method with 1,000 bootstrap replicates under the Poisson correction model. Sequence alignments were visualized with ESPript 3.0 (https://espript.ibcp.fr/ESPript/cgi-bin/ESPript.cgi).

### Protein structure prediction and homology identification

The three-dimensional structure of *A. thaliana* HWS was predicted by AlphaFold3 (Abramson et al., 2024) and visualized in PyMOL (The PyMOL Molecular Graphics System, Schrödinger, LLC).

### AGO1-RNA Immunoprecipitation

AGO1-RIP analysis was performed following established protocols (Mei et al., 2019). Approximately 3 mL of finely ground seedling tissue was prepared in liquid nitrogen and homogenized in 5 mL of IP buffer [30 mM Tris–HCl (pH 7.5), 5 mM MgCl₂, 150 mM NaCl, 80 mM potassium acetate, 5% glycerol, 0.5% Triton X-100, 2.5 mM DTT, 0.035% β-mercaptoethanol, 1 mM PMSF, 40 U/mL RNase inhibitor, and protease inhibitor cocktail (Roche, No.11873580001)]. After removing the debris, the supernatant was incubated with 35 μL of anti-AGO1-conjugated protein A/G agarose beads at 4°C for 4 h. A 5% aliquot of the beads was reserved for AGO1 detection by western blotting, while the remaining beads were eluted with RIP elution buffer [100 mM Tris–HCl (pH 7.5), 10 mM EDTA (pH 8.0), 1% SDS, 50 U/mL RNase inhibitor] at 25°C for 10 min. RNAs from the eluted fractions were extracted with phenol-chloroform (1:1, V/V), desalted using a G-25 column (Cytiva), and precipitated by adding three volumes of ethanol at −80°C for 3 h. The final RNA pellet was dissolved in 15 μL DEPC-treated water. Of this, 5 μL was used for PCR analysis, and the remaining 10 μL was used for Northern blot.

## Acknowledgements

We are grateful to Dr. Binglian Zheng for sharing the custom-made anti-AGO1 antibody. This work was supported by Shanghai Pilot Program for Basic Research-Fudan University 21TQ1400100 (22TQ014), Science and Technology Commission of Shanghai Municipality (22ZR1406100), and the National Natural Science Foundation of China (32470591).

## Author contributions

G.R., L.Y., and J.M. designed the study. L.Y., M.G., Y. Fan., and Y. Fang. performed the experiments. M.L., G.K., and J.Y. designed and generated the prime editing toolkit. L.Y., N.L., Y.W., X.Z., and G.R. analyzed the data. L.Y. and G.R. wrote the manuscript.

## Reference

Abe A, Kosugi S, Yoshida K, Natsume S, Takagi H, Kanzaki H, Matsumura H, Yoshida K, Mitsuoka C, Tamiru M et al (2012) Genome sequencing reveals agronomically important loci in rice using MutMap. Nat Biotechnol 30: 174–178

Abramson J, Adler J, Dunger J, Evans R, Green T, Pritzel A, Ronneberger O, Willmore L, Ballard AJ, Bambrick J et al (2024) Accurate structure prediction of biomolecular interactions with AlphaFold 3. Nature 630: 493–500

Ahmad A, Cao X (2012) Plant PRMTs broaden the scope of arginine methylation. J Genet Genomics 39: 195–208

Barre-Villeneuve C, Laudie M, Carpentier MC, Kuhn L, Lagrange T, Azevedo-Favory J (2024) The unique dual targeting of AGO1 by two types of PRMT enzymes promotes phasiRNA loading in *Arabidopsis* thaliana. Nucleic Acids Res 52: 2480–2497

Borah P, Khurana JP (2018) The OsFBK1 E3 Ligase Subunit Affects Anther and Root Secondary Cell Wall Thickenings by Mediating Turnover of a Cinnamoyl-CoA Reductase. Plant Physiol 176: 2148–2165

Borna RS, Murchie EH, Pyke KA, Roberts JA, Gonzalez-Carranza ZH (2022) The rice EP3 and OsFBK1 E3 ligases alter plant architecture and flower development, and affect transcript accumulation of microRNA pathway genes and their targets. Plant Biotechnol J 20: 297–309

Chen J, Liu L, You C, Gu J, Ruan W, Zhang L, Gan J, Cao C, Huang Y, Chen X et al (2018) Structural and biochemical insights into small RNA 3’ end trimming by *Arabidopsis* SDN1. Nat Commun 9: 3585

Damayanti F, Lombardo F, Masuda JI, Shinozaki Y, Ichino T, Hoshikawa K, Okabe Y, Wang N, Fukuda N, Ariizumi T et al (2019) Functional Disruption of the Tomato Putative Ortholog of HAWAIIAN SKIRT Results in Facultative Parthenocarpy, Reduced Fertility and Leaf Morphological Defects. Front Plant Sci 10: 1234

Franco-Zorrilla JM, Valli A, Todesco M, Mateos I, Puga MI, Rubio-Somoza I, Leyva A, Weigel D, Garcia JA, Paz-Ares J (2007) Target mimicry provides a new mechanism for regulation of microRNA activity. Nat Genet 39: 1033–1037

Gonzalez-Carranza ZH, Rompa U, Peters JL, Bhatt AM, Wagstaff C, Stead AD, Roberts JA (2007) Hawaiian skirt: an F-box gene that regulates organ fusion and growth in *Arabidopsis*. Plant Physiol 144: 1370–1382

Gonzalo L, Gagliardi D, Zlauvinen C, Gulanicz T, Arce AL, Fernandez J, Cambiagno DA, Merchante C, Zienkiewicz A, Jarmolowski A et al (2025) The nuclear pore complex acts as a hub for pri-miRNA transcription and processing in plants. Nucleic Acids Res 53: gkaf885

Guan S, Tan SM, Li Y, Torres J, Alex Law SK (2016) Function and conformation analyses of an aspartate substitution of the invariant glycine in the integrin betaI domain alpha1-alpha1’ helix. Biochem Biophys Rep 7: 214–217

Han J, LaVigne CA, Jones BT, Zhang H, Gillett F, Mendell JT (2020) A ubiquitin ligase mediates target-directed microRNA decay independently of tailing and trimming. Science 370: eabc9546

Hassan Mu, Zainal Z, Ismail I (2015) Plants kelch containing F-box proteins-Structure, Evolution and Functions. RCS Advances 5: 42808–42814

Hu P, Zhao H, Zhu P, Xiao Y, Miao W, Wang Y, Jin H (2019) Dual regulation of *Arabidopsis* AGO2 by arginine methylation. Nat Commun 10: 844

Ito N, Phillips S, Stevens C, Ogel Z, McPherson M, Keen J, Yadav K, Knowles P (1991) Novel thioether bond revealed by a 1.7 Å crystal structure of galactose oxidase. Nature 350: 87–90

Lang PLM, Christie MD, Dogan ES, Schwab R, Hagmann J, van de Weyer AL, Scacchi E, Weigel D (2018) A Role for the F-Box Protein HAWAIIAN SKIRT in Plant microRNA Function. Plant Physiol 176: 730–741

Li J, Yang Z, Yu B, Liu J, Chen X (2005) Methylation protects miRNAs and siRNAs from a 3’-end uridylation activity in *Arabidopsis*. Curr Biol 15: 1501–1507

Li M, Tang D, Wang K, Wu X, Lu L, Yu H, Gu M, Yan C, Cheng Z (2011) Mutations in the F-box gene LARGER PANICLE improve the panicle architecture and enhance the grain yield in rice. Plant Biotechnol J 9: 1002–1013

Li M, Yu B (2021) Recent advances in the regulation of plant miRNA biogenesis. RNA Biol 18: 2087–2096

Li S, Castillo-Gonzalez C, Yu B, Zhang X (2017) The functions of plant small RNAs in development and in stress responses. Plant J 90: 654–670

Li X, Zhang D, Hannink M, Beamer LJ (2004) Crystal structure of the Kelch domain of human Keap1. J Biol Chem 279: 54750–54758

Liu Y, Nakatsukasa K, Kotera M, Kanada A, Nishimura T, Kishi T, Mimura S, Kamura T (2011) Non-SCF-type F-box protein Roy1/Ymr258c interacts with a Rab5-like GTPase Ypt52 and inhibits Ypt52 function. Mol Biol Cell 22: 1575–1584

Lombardo F, Gramazio P, Ezura H (2021) Increase in Phloem Area in the Tomato hawaiian skirt Mutant Is Associated with Enhanced Sugar Transport. Genes (Basel) 12: 932

Mei J, Jiang N, Ren G (2019) The F-box protein HAWAIIAN SKIRT is required for mimicry target-induced micoRNA degradation. J Integr Plant Biol 61: 1121–1127

Ogura Y, Ihara N, Komatsu A, Tokioka Y, Nishioka M, Takase T, Kiyosue T (2008) Gene expression, localization, and protein–protein interaction of *Arabidopsis* SKP1-like (ASK) 20A and 20B. Plant Science 174: 485–495

Peng T, Qiao M, Liu H, Teotia S, Zhang Z, Zhao Y, Wang B, Zhao D, Shi L, Zhang C et al (2018) A Resource for Inactivation of MicroRNAs Using Short Tandem Target Mimic Technology in Model and Crop Plants. Mol Plant 11: 1400–1417

Piao R, Jiang W, Ham TH, Choi MS, Qiao Y, Chu SH, Park JH, Woo MO, Jin Z, An G et al (2009) Map-based cloning of the ERECT PANICLE 3 gene in rice. Theor Appl Genet 119: 1497–1506

Ramachandran V, Chen X (2008) Degradation of microRNAs by a family of exoribonucleases in *Arabidopsis*. Science 321: 1490–1492

Rapp C, Klerman H, Levine E, McClendon CL (2013) Hydrogen bond strengths in phosphorylated and sulfated amino acid residues. PLoS One 8: e57804

Reichel M, Li Y, Li J, Millar AA (2015) Inhibiting plant microRNA activity: molecular SPONGEs, target MIMICs and STTMs all display variable efficacies against target microRNAs. Plant Biotechnol J 13: 915–926

Ren G, Chen X, Yu B (2012) Uridylation of miRNAs by hen1 suppressor1 in *Arabidopsis*. Curr Biol 22: 695–700

Ren G, Xie M, Zhang S, Vinovskis C, Chen X, Yu B (2014) Methylation protects microRNAs from an AGO1-associated activity that uridylates 5’ RNA fragments generated by AGO1 cleavage. Proc Natl Acad Sci U S A 111: 6365–6370

Rieu P, Arnoux-Courseaux M, Tichtinsky G, Parcy F (2023a) Thinking outside the F-box: how UFO controls angiosperm development. New Phytol 240: 945–959

Rieu P, Turchi L, Thévenon E, Zarkadas E, Nanao M, Chahtane H, Tichtinsky G, Lucas J, Blanc-Mathieu R, Zubieta C et al (2023b) The F-box protein UFO controls flower development by redirecting the master transcription factor LEAFY to new cis-elements. Nature Plants 9: 315–329

Sheu-Gruttadauria J, Pawlica P, Klum SM, Wang S, Yario TA, Schirle Oakdale NT, Steitz JA, MacRae IJ (2019) Structural Basis for Target-Directed MicroRNA Degradation. Mol Cell 75: 1243–1255

Shi CY, Kingston ER, Kleaveland B, Lin DH, Stubna MW, Bartel DP (2020) The ZSWIM8 ubiquitin ligase mediates target-directed microRNA degradation. Science 370: eabc9359

Shin H, Shin HS, Chen R, Harrison MJ (2006) Loss of At4 function impacts phosphate distribution between the roots and the shoots during phosphate starvation. The Plant Journal 45: 712–726

Song X, Li Y, Cao X, Qi Y (2019) MicroRNAs and Their Regulatory Roles in Plant-Environment Interactions. Annu Rev Plant Biol 70: 489–525

Su L-Y, Li S-S, Liu H, Cheng Z-M, Xiong A-S (2024) The origin, evolution, and functional divergence of the Dicer-like (DCL) and Argonaute (AGO) gene families in plants. Epigenetics Insights 17: 0–0

Todesco M, Rubio-Somoza I, Paz-Ares J, Weigel D (2010) A collection of target mimics for comprehensive analysis of microRNA function in *Arabidopsis thaliana*. PLoS Genet 6: e1001031

Tu B, Liu L, Xu C, Zhai J, Li S, Lopez MA, Zhao Y, Yu Y, Ramachandran V, Ren G et al (2015) Distinct and cooperative activities of HESO1 and URT1 nucleotidyl transferases in microRNA turnover in *Arabidopsis*. PLoS Genet 11: e1005119

Vaucheret H, Vazquez F, Crete P, Bartel DP (2004) The action of ARGONAUTE1 in the miRNA pathway and its regulation by the miRNA pathway are crucial for plant development. Genes Dev 18: 1187–1197

Vierstra RD (2009) The ubiquitin-26S proteasome system at the nexus of plant biology. Nat Rev Mol Cell Biol 10: 385–397

Wang J, Mei J, Ren G (2019) Plant microRNAs: Biogenesis, Homeostasis, and Degradation. Front Plant Sci 10: 360

Wang X, Wang Y, Dou Y, Chen L, Wang J, Jiang N, Guo C, Yao Q, Wang C, Liu L et al (2018) Degradation of unmethylated miRNA/miRNA*s by a DEDDy-type 3’ to 5’ exoribonuclease Atrimmer 2 in *Arabidopsis*. Proc Natl Acad Sci U S A 115: E6659–E6667

Wang X, Zhang S, Dou Y, Zhang C, Chen X, Yu B, Ren G (2015) Synergistic and independent actions of multiple terminal nucleotidyl transferases in the 3’ tailing of small RNAs in *Arabidopsis*. PLoS Genet 11: e1005091

Xu W, Yang Y, Yang B, Krueger CJ, Xiao Q, Zhao S, Zhang L, Kang G, Wang F, Yi H et al (2022) A design optimized prime editor with expanded scope and capability in plants. Nature Plants 8: 45–52

Xu Y, Chen X (2023) microRNA biogenesis and stabilization in plants. Fundam Res 3: 707–717

Yan J, Gu Y, Jia X, Kang W, Pan S, Tang X, Chen X, Tang G (2012) Effective small RNA destruction by the expression of a short tandem target mimic in *Arabidopsis*. Plant Cell 24: 415–427

Yang L, Li N, Guo M, Mei J, Ren G (2022) HWS as a central player in target mimicry induced miRNA degradation. Journal of Fudan Univerity ( Natural Science) 61: 375–384

Yang Y, Hadjikyriacou A, Xia Z, Gayatri S, Kim D, Zurita-Lopez C, Kelly R, Guo A, Li W, Clarke SG et al (2015) PRMT9 is a type II methyltransferase that methylates the splicing factor SAP145. Nat Commun 6: 6428

Yu B, Yang Z, Li J, Minakhina S, Yang M, Padgett R, Steward R, Chen X (2005) Methylation as a crucial step in plant. Science 307: 932–935

Yu H, Murchie EH, Gonzalez-Carranza ZH, Pyke KA, Roberts JA (2015) Decreased photosynthesis in the erect panicle 3 (ep3) mutant of rice is associated with reduced stomatal conductance and attenuated guard cell development. J Exp Bot 66: 1543–1552

Yu Y, Ji L, Le BH, Zhai J, Chen J, Luscher E, Gao L, Liu C, Cao X, Mo B et al (2017) ARGONAUTE10 promotes the degradation of miR165/6 through the SDN1 and SDN2 exonucleases in *Arabidopsis*. PLoS Biol 15: e2001272

Yu Y, Wang H, You C, Chen X (2025) Plant microRNA maturation and function. Nat Rev Mol Cell Biol

Zhai J, Zhao Y, Simon SA, Huang S, Petsch K, Arikit S, Pillay M, Ji L, Xie M, Cao X et al (2013) Plant microRNAs display differential 3’ truncation and tailing modifications that are ARGONAUTE1 dependent and conserved across species. Plant Cell 25: 2417–2428

Zhang H, Zhang J, Yan J, Gou F, Mao Y, Tang G, Botella JR, Zhu JK (2017a) Short tandem target mimic rice lines uncover functions of miRNAs in regulating important agronomic traits. Proc Natl Acad Sci U S A 114: 5277–5282

Zhang X, Gonzalez-Carranza ZH, Zhang S, Miao Y, Liu CJ, Roberts JA (2019) F-Box Proteins in Plants. In: Annual Plant Reviews pp. 307–328.

Zhang X, Jayaweera D, Peters JL, Szecsi J, Bendahmane M, Roberts JA, Gonzalez-Carranza ZH (2017b) The *Arabidopsis thaliana* F-box gene HAWAIIAN SKIRT is a new player in the microRNA pathway. PLoS One 12: e0189788

Zhao Y, Yu Y, Zhai J, Ramachandran V, Dinh TT, Meyers BC, Mo B, Chen X (2012) The *Arabidopsis* nucleotidyl transferase HESO1 uridylates unmethylated small RNAs to trigger their degradation. Curr Biol 22: 689–694

Zuber H, Scheer H, Joly AC, Gagliardi D (2018) Respective Contributions of URT1 and HESO1 to the Uridylation of 5’ Fragments Produced From RISC-Cleaved mRNAs. Front Plant Sci 9: 1438

