## Supplemental Figures 1-4 for "An extensive forward genetic screen identifies the HWS-AGO1 axis as pivotal for target mimicry-mediated miRNA degradation in *Arabidopsis*"

### Supplemental Figures and Legends

**A**

| Allele | Genomic mutation | AA Substitution | Domain | Reference |
|---|---|---|---|---|
| *hws-1* | A 28bp deletion resulted in a premature translation termination | | Kelch-Repeat_5 | González-Carranza et al., 2007. |
| *hws-2* | T-DNA insertion (SALK_088349) | | Kelch-Repeat_2 | González-Carranza et al., 2007. |
| *hws-3 shr* | C516A | Y172* | Kelch-Repeat_3 | Kim et al., 2016. |
| *hws-4 shr* | G993A | W331* | Kelch-Repeat_5 | Kim et al., 2016. |
| *hws-5 MIM156* | G537A | W179* | Kelch-Repeat_3 | Lang et al.,2018. |
| *hws-6 STTM160* | G345A | W115* | Kelch-Repeat_1 | Mei et al.,2018. |
| *hws-7 STTM160* | G920A | G307E | Kelch-Repeat_5 | Mei et al.,2018. |
| *hws-8 STTM160* | G1067A | G356D | Kelch-Repeat_6 | Yang et al.,2021. |
| *hws-9 STTM160* | C167T | P56L | F-Box | Yang et al.,2021. |
| *hws-10 STTM160* | G705A | W235* | Kelch-Repeat_4 | Yang et al.,2021. |
| *hws-11 STTM160* | G314A | G105D | Kelch-Repeat_1 | Yang et al.,2021. |
| *hws-12 STTM160* | G1199A | G400E | Kelch-Repeat_1 | Yang et al.,2021. |
| *hws-13 STTM160* | G1039A | E347K | Kelch-Repeat_6 | Yang et al.,2021. |
| *hws-14 STTM160* | G209A | W70* | F-Box | Yang et al.,2021. |
| *hws-15 STTM160* | G718A | E240K | Kelch-Repeat_4 | This work |
| *hws-16 STTM160* | G1048A | E350K | Kelch-Repeat_6 | This work |
| *hws-17 STTM160* | G1149A | W383* | Kelch-Repeat_6 | This work |
| *hws-18 STTM160* | G191A | G64N | F-Box | This work |
| *hws-19 STTM160* | G384A | W128* | Kelch-Repeat_2 | This work |
| *hws-20 STTM160* | G630A | W210* | Kelch-Repeat_3 | This work |
| *hws-21 STTM160* | G972A | W324* | Kelch-Repeat_5 | This work |
| *hws-22 STTM160* | G1061A | S354N | Kelch-Repeat_6 | This work |
| *hws-23 STTM160* | G1099A | G367R | Kelch-Repeat_6 | This work |
| *hws-24 STTM160* | G971A | W324* | Kelch-Repeat_5 | This work |
| *hws-25 STTM160* | C1015T | Q339* | Kelch-Repeat_6 | This work |
| *hws-26 STTM160* | C1013T | P338L | Kelch-Repeat_5 | This work |
| *hws-27 STTM160* | G1143A | W381* | Kelch-Repeat_6 | This work |
| *hws-28 STTM160* | G1100A | G367E | Kelch-Repeat_6 | This work |

Note: *hws-21* and *hws-24* carried the identical *HWS* mutation, resulting in a premature stop codon at residue 324.

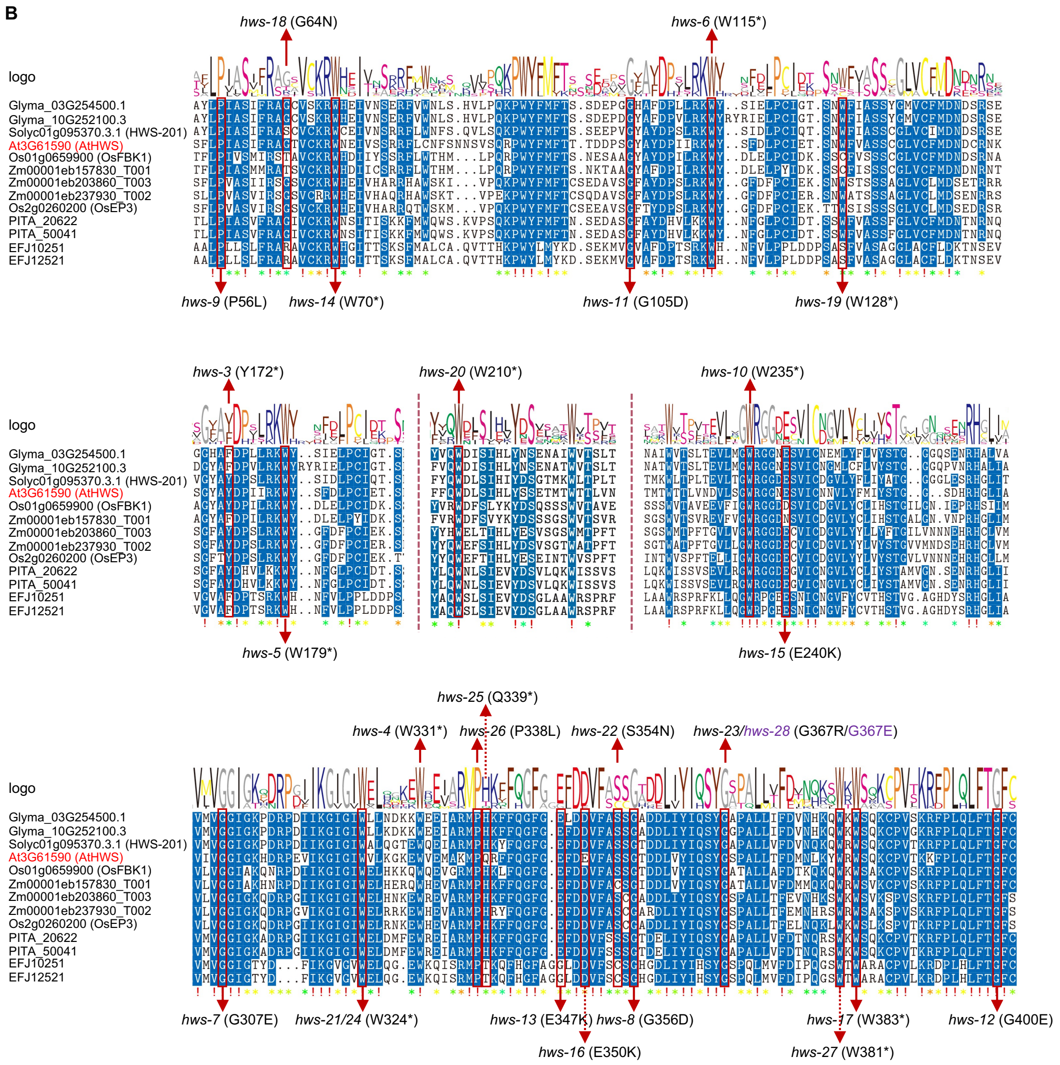


**Fig. S1.** *hws* allelic mutations. **(A)** Summary of known and newly identified *hws* mutants. **(B)** Conservation of the mutated residues among HWS homologs across plant species, including *Arabidopsis thaliana* (At), *Glycine max* (Glyma), *Solanum lycopersicum* (Solyc), *Oryza sativa* (Os), *Zea mays* (Zm), *Pinus taeda* (PITA), and *Selaginella moellendorffii* (EFJ).

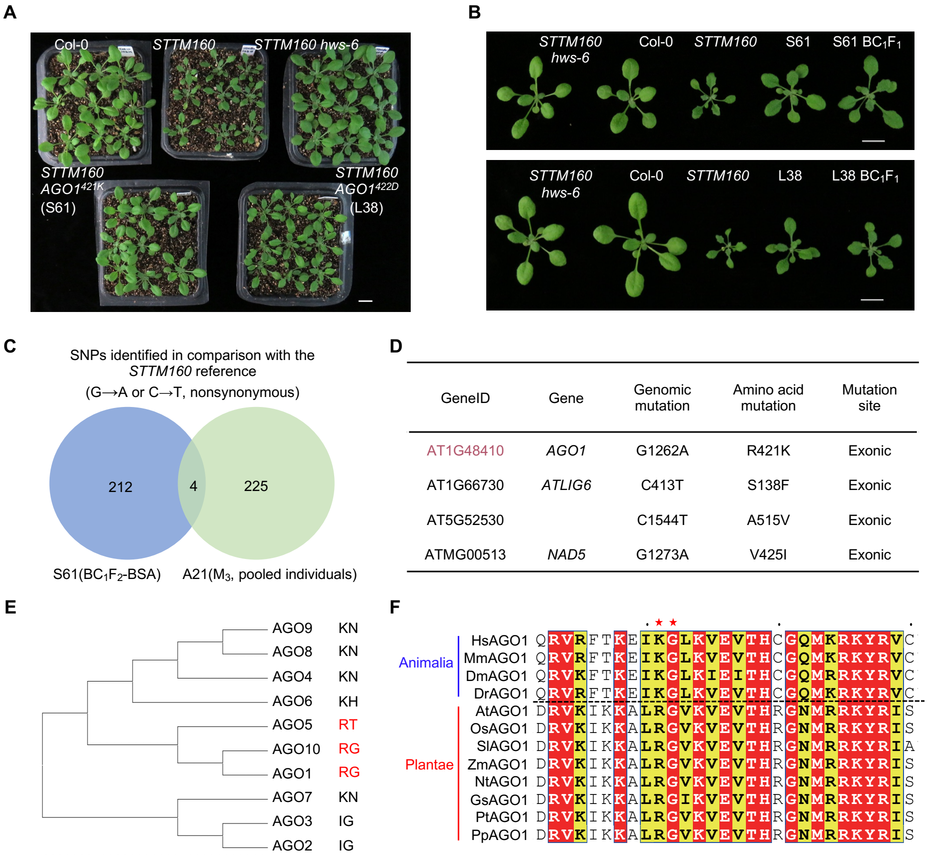


**Fig. S2.** Characterization of the *STTM160 AGO1*$^{K421}$ and *STTM160 AGO1*$^{D422}$ mutants. **(A)** Three-week-old plants of the indicated genotypes. Scale bar = 1 cm. **(B)** Backcross analysis. S61BC$_1$F$_1$, the F$_1$ progeny from the backcross between S61 (*STTM160 AGO1*$^{K421}$) and *STTM160*. L38BC$_1$F$_1$, the F1 progeny from the backcross between L38 (*STTM160 AGO1*$^{D422}$) and *STTM160*. Morphological phenotypes of three-week-old plants were shown. Scale bar = 1 cm. **(C&D)** A21 is an independent suppressor line carrying the same AGO1 mutation (c.1262G>A) as S61. (**C**) A Venn diagram showing overlaps of detected SNPs. **(D)** The four shared SNPs between S61 and A21 include the mutation in *AGO1*. **(E)** Conservation analysis of the AGO1 R421 and G422 residues. Species analyzed are *Homo sapiens* (Hs), *Mus musculus* (Mm), *Drosophila melanogaster* (Dm), *Danio rerio* (Dr), *Oryza sativa subsp. Japonica* (Os), *Solanum lycopersicum* (Sl), *Populus tomentosa* (Pt), *Zea mays* (Zm), *Physcomitrella patens* (Pp), *Nicotiana tabacum* (Nt), *Glycine soja* (Gs).

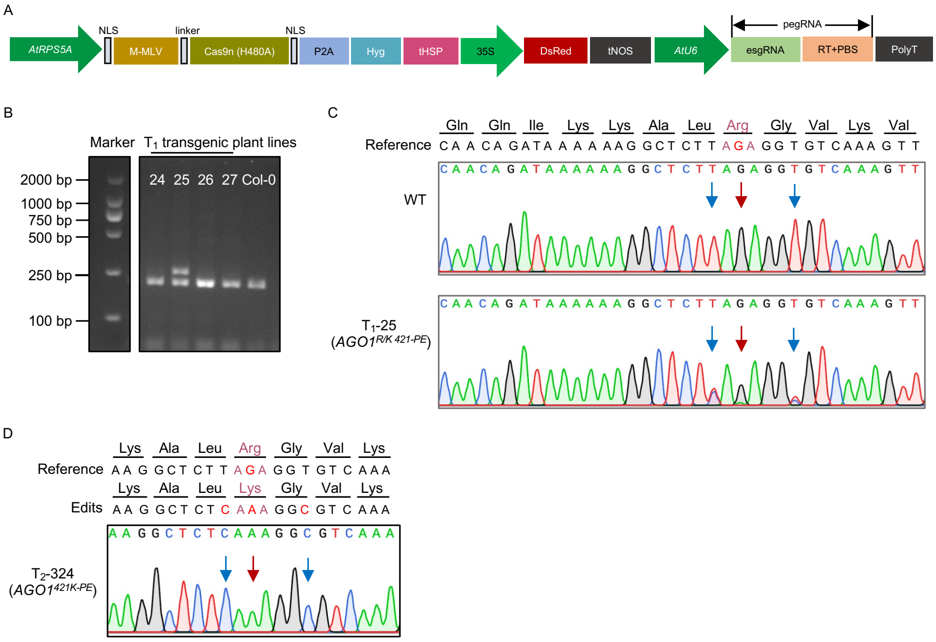


**Fig. S3.** Generation of *AGO1*$^{421K}$-edited plant by a prime editing system. **(A)** Schematic illustration of the PE construct. P2A, self-cleaving peptide 2A; esgRNA, enhanced small guide RNA; pegRNA, prime editing guide RNA; tHSP, terminator of *AtHSP18.2*. **(B)** Representative dCAPS-based genotyping of independent $T_1$ transformants. PCR amplicons were digested with DdeI for 12 h and resolved by agarose gel electrophoresis. **(C)** Sanger sequencing chromatogram of a representative $T_1$-edited plant. The red arrow marks the target editing site, and the blue arrowhead denotes a synonymous substitution introduced in the reverse transcription template to facilitate editing efficiency. **(D)** Sanger sequencing chromatogram of a homozygous edited line isolated from the $T_2$ generation.

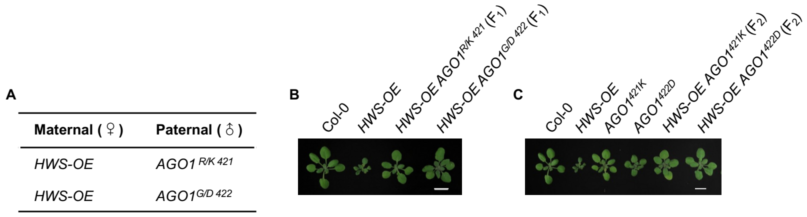


**Fig. S4.** *AGO1*$^{K421}$ and *AGO1*$^{D422}$ suppressed the developmental defects of HWS overexpression (*HWS-OE*) in a dominant manner. **(A)** Parental lines used for crosses. For *HWS-OE*, a single-copied transgenic line with stable developmental defects was used. **(B)** Vegetative phenotypes of $F_1$ plants with indicated genotypes. **(C)** Vegetative phenotypes of $F_2$ plants with indicated genotypes. Aerial parts of three-week-old plants were photographed. Scale bar = 1 cm.

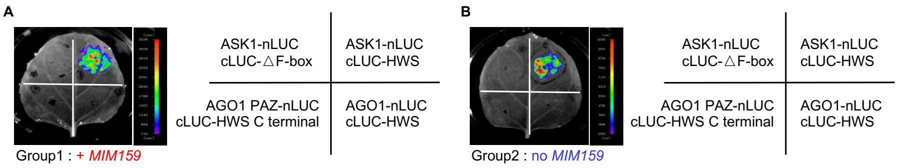


**Fig. S5.** Split-luciferase analysis of the interaction between AGO1 and HWS with **(A)** or without **(B)** co-infiltration of *MIM159*. Indicated protein pairs (text in the right panel) were infiltrated in tobacco leaves, and luminescence signals were detected after 48 hours.

#### Supplemental Table:

Table S1 Primers and oligos used in this study.

#### References:

Gonzalez-Carranza ZH, Rompa U, Peters JL, Bhatt AM, Wagstaff C, Stead AD, Roberts JA (2007) Hawaiian skirt: an F-box gene that regulates organ fusion and growth in *Arabidopsis*. *Plant Physiol* 144: 1370-1382

Kim ES, Choe G, Sebastian J, Ryu KH, Mao L, Fei Z, Lee JY (2016) HAWAIIAN SKIRT regulates the quiescent center-independent meristem activity in *Arabidopsis* roots. *Physiol Plant* 157: 221-233

Lang PLM, Christie MD, Dogan ES, Schwab R, Hagmann J, van de Weyer AL, Scacchi E, Weigel D (2018) A Role for the F-Box Protein HAWAIIAN SKIRT in Plant microRNA Function. *Plant Physiol* 176: 730-741

Mei J, Jiang N, Ren G (2019) The F-box protein HAWAIIAN SKIRT is required for mimicry target-induced microRNA degradation. *J Integr Plant Biol* 61: 1121-1127

Yang L, Li N, Guo M, Mei J, Ren G (2022) HWS as a central player in target mimicry induced miRNA degradation. *Journal of Fudan Univerity (Natural Science)* 61: 375-384
